# The life-history fitness of F_1_ hybrids of the microcrustacean *Daphnia pulex* and *D. pulicaria* (Crustacea, Anomopoda)

**DOI:** 10.1101/2020.08.28.271791

**Authors:** Irene Moy, Makayla Green, Thinh Phu Pham, Dustin Luu, Sen Xu

**Affiliations:** Department of Biology, University of Texas at Arlington, Texas, 76019, USA

**Author notes:** Correspondence: 501 S. Nedderman Dr, Arlington, TX, 76019.

**Keywords:** heterosis, speciation, genetic incompatibility, hybridization, life history

## Abstract

Negative interaction between alleles that arise independently in diverging populations (i.e., Dobzhansky-Muller incompatibilities) can cause reduction of fitness in their hybrids. However, heterosis in hybrids can emerge if hybridization breaks down detrimental epistatic interaction within parental lineages. In this study, we examined the life-history fitness of the inter-specific F_1_ hybrids of two recently diverged microcrustacean species *Daphnia pulex* and *D. pulicaria* as well as intra-specific F_1_ hybrids of *D. pulex*. We identified heterosis in four out of five life-history traits in the inter-specific F1 hybrids. The observation of heterosis in these life-history traits suggests that there are no major genetic incompatibilities between these two species affecting these traits. This also suggests that *D. pulex* and *D. pulicaria* are at the early stage of speciation where heterosis can still occur.

## Introduction

Evolutionary theory predicts that a major step in speciation is the reduction of fitness of hybrids between two diverging lineages (Coyne & Orr, 2004), which can reinforce the divergence of differentiating lineages. The reduction of fitness in hybrids is usually attributed to the Dobzhansky-Muller incompatibilities (Dobzhansky, 1937; Muller, 1940), which are negative interaction between alleles that arise in the diverging populations and are brought together in hybrids for the first time. Over the years much empirical evidence supporting the Dobzhansky-Muller incompatibilities has emerged (Barbash et al., 2003; Mack & Nachman, 2016; Presgraves, 2003; Rawson & Burton, 2002). It is predicted that with longer divergence time between isolated populations the number of accumulated incompatibilities quadratically increases, eventually leading to hybrid lethality and sterility (Coyne & Orr, 2004).

However, there are also many cases of recently diverged lineages where hybrids are more fit than either parents, i.e., heterosis (Bernardes et al., 2017; Birchler et al., 2010; Bolnick & Near, 2005; Edmands, 1999). The heterosis in hybrids can be explained by uniparentally inherited elements (Fraisse et al., 2016) or low fitness in parents (Simon et al., 2018). Interestingly, recent theories predict that heterosis can arise transiently during the early stage of speciation process if the epistatic interaction in the parental populations is deleterious and the disruption of these deleterious interaction in the F_1_ hybrids can result in elevated fitness (Dagilis et al., 2019).

Given these empirical data and theoretical insights into hybrid fitness for diverging lineages, we set out to evaluate the fitness of F_1_ hybrids between two recently diverged microcrustacean species *Daphnia pulex* and *D. pulicaria* in North America. *Daphnia pulex* and *D. pulicaria* are member species of the *D. pulex* species complex (Colbourne et al., 1998; Colbourne & Hebert, 1996). *Daphnia* is well known for its cyclically parthenogenetic life cycle. Under favorable environmental conditions (e.g., ample food and low population density), populations mainly consist of females and females reproduce asexually to produce genetically identical daughters for several generations (Ebert, 2005). When environmental conditions deteriorate (e.g., low food condition and high population density), males will appear in the population from the asexually produced neonates as sex is environmentally determined in *Daphnia*. Meantime, females can perceive these environmental cues and switch to sexual reproduction to generate haploid egg through meiosis, which upon fertilization by sperm becomes fertilized resting embryos. The sexually produced resting embryos can survive in harsh environmental conditions as they are encapsulated in a protective structure (i.e., ephippium), and they can hatch out once the environmental conditions become suitable again (Ebert, 2005).

While *D. pulex* and *D. pulicaria* diverged about 1-2 million years ago (Cristescu et al., 2012; Omilian & Lynch, 2009), there is little morphological divergence between these two species (Brandlova et al., 1972). However, they inhabit distinct habitats in overlapping geographic regions. *D. pulex* exclusively lives in fishless ephemeral ponds and *D. pulicaria* occupying stratified permanent lakes (Cristescu et al., 2012). Despite their recent divergence, these two species most likely underwent local adaptation as they show distinct life history and physiological traits. For example, *D. pulicaria* has a diel vertical migration behavior, coming up to the surface of lake during night and staying in deep water column during the day, whereas *D. pulex* living in ponds does not have such behavior. Notably, prezygotic isolation has developed between these two species (Deng 1997), with *D. pulex* switching to sexual reproduction at long-day hours (16 hours/day) and *D. pulicaria* switching to sexual at short-day hours (10 hours/day).

Although *D. pulex* and *D. pulicaria* can generate fertile cyclically parthenogenetic F_1_ offspring in laboratory crossing experiments (Heier and Dudycha 2009), the *D. pulex*-*D. pulicaria* hybrid populations that are found in the field so far are mostly obligately parthenogenetic (Hebert et al., 1989), i.e., they lost the capability of sexual reproduction and produces ephippial resting embryos through parthenogenesis rather than by sex. The genetic mechanisms underlying this switch in reproduction mode of hybrid populations is an active research area (Tucker et al., 2013; Xu et al., 2013; Xu et al., 2015). It is noted that the recorded hybrid populations are of *D. pulex* maternal ancestry and *D. pulicaria* paternal ancestry based on mitochondrial DNA genealogy, suggesting unidirectional hybridization between these two species in nature (Chin et al., 2019; Cristescu et al., 2012; Heier & Dudycha, 2009).

In this study, we specifically examine whether F_1_ hybrids between *D. pulex* and *D. pulicaria* show heterosis (i.e., higher fitness than either parents). We performed crossing experiments between males of *D. pulicaria* and females of *D. pulex* to generate F_1_s. As *Daphnia* mainly exist as females during their life history, we evaluated the female life history traits in the F_1_s and parental lineages. We also performed crosses between *D. pulex* populations to assess whether the intra-specific outbred hybrids resemble the life-history patterns of inter-specific hybrids.

## Materials and Methods

### Crossing experiments

We performed two inter-specific crossing experiment and one intra-specific crossing experiment. For the inter-specific crossing, we crossed males of a *D. pulicaria* genotype AroMoose with females of *D. pulex* genotypes PA32 and Tex21, respectively, whereas for the intra-specific crossing males of *D. pulex* genotype Tex21 were crossed with females of *D. pulex* genotype PA32. To perform crossing experiments, 20 males of the paternal genotype and 20 females of the maternal genotype were placed together in a 250 mL beaker at 18 °C under 12:12 (Light:Dark) photoperiod. We stressed the animals with low water volume and low food conditions to stimulate the females to switch to sexual reproduction. Asexually produced babies were removed every 2 days to avoid crossing between males and females of the same genotype. Sexually produced embryos contained in ephippia were collected every 2 days and dissected. Collected embryos were then stored in the dark for 2-3 weeks and exposed to 410 nm UV light to stimulate hatching as described in Luu et al., (2020). The hatched neonates were collected and separately maintained in lake combo and 12:12 (L:D) photoperiod at 18 °C to establish clonal lines.

### Life history assay

We measured five life-history traits on the parental genotypes and F_1_ hybrids in common garden experiments. These traits are body size of females at sexual maturity, number of days to reach female sexual maturity, number of asexual broods during life span, brood size, and body size of 2-3 old neonates.

To avoid maternal effects on these life history traits, asexual females of all examined parental and F_1_ genotypes were raised for two generations in artificial lake water COMBO (Kilham et al., 1998) under 12:12 (L:D) photoperiod at 18 °C. We fed the animals with the green algae *Scenedesmus obliquus* at a concentration of 500,000 cells per ml and changed the artificial lake water every two days. When the second generation of each *Daphnia* isolate started reproducing asexually, we randomly selected at least 15 female babies, kept each of them separately at the standard culture conditions, and performed life-history trait assays.

### Number of days to maturity

The number of days to maturity was calculated as the number of days between the birth date and the date for the first sighting of asexual embryos deposited in the brood pouch for each focal individual.

### Body size of females at sexual maturity

The body size of each individual was measured on the day of first sighting of asexual embryos in the brood pouch. A picture was taken using the Leica M125 in the computer program Leica Application Suite V4. We measured the distance between the top of the head to the bottom of the carapace in the picture. The tail spine was not included in the measurements as it often differed in size or were deformed.

### Number of asexual broods during life span and brood size

After noting that individuals reached sexual maturity, we continued to observe individual *Daphnia* every day and recorded the number of asexual broods and brood sizes that each individual had until it died. All asexually produced neonates were removed from the container upon their birth to keep standardized environmental conditions for each individual. These neonates were measured for their body sizes as described below.

### Body size of 3-day old neonates

Asexually produced babies were placed in a container under standardized conditions until they reached 3 days old. It is noted that each brood of neonates were placed together in a separate container. Once the neonates reached 3 day old, they were measured for body size. For each G_3_ *Daphnia* individual, a total of 3 broods were measured. If the brood size was ≤ 10 babies, all neonates were measured. If the brood size was larger than 10 babies, 10 babies were randomly selected and measured. The body size was calculated as the distance between the top of the head to the bottom of carapace. The tail spine was not included in the measurements as it often differed in size or was deformed.

## Results

We identified the parental strains differed in multiple life-history traits that we examined. Cases of heterosis were identified in four out of the five life-history traits, whereas hybrid depression was observed at a lower frequency. The mean and standard deviation were presented for all the traits below. The p values in this section were all derived from t-test unless otherwise noted.

### Female body size at maturity

Among the parental isolates used for crossing experiments, we found that the average female body size was 1.65 ±0.06 (SD) mm for PA32 (*D. pulex*), 1.65 ±0.12 mm for Tex21 (*D. pulex*), and 1.82 ±0.10 mm for AroMoose (*D. pulicaria*). Pairwise t-test showed that AroMoose (*D. pulicaria*) has a significantly bigger body size than either *D. pulex* isolates PA32 or Tex21 (*p* < 0.001, **Figure 1**), whereas no significant difference was detected between two *D. pulex* isolates.

**Figure 1.**
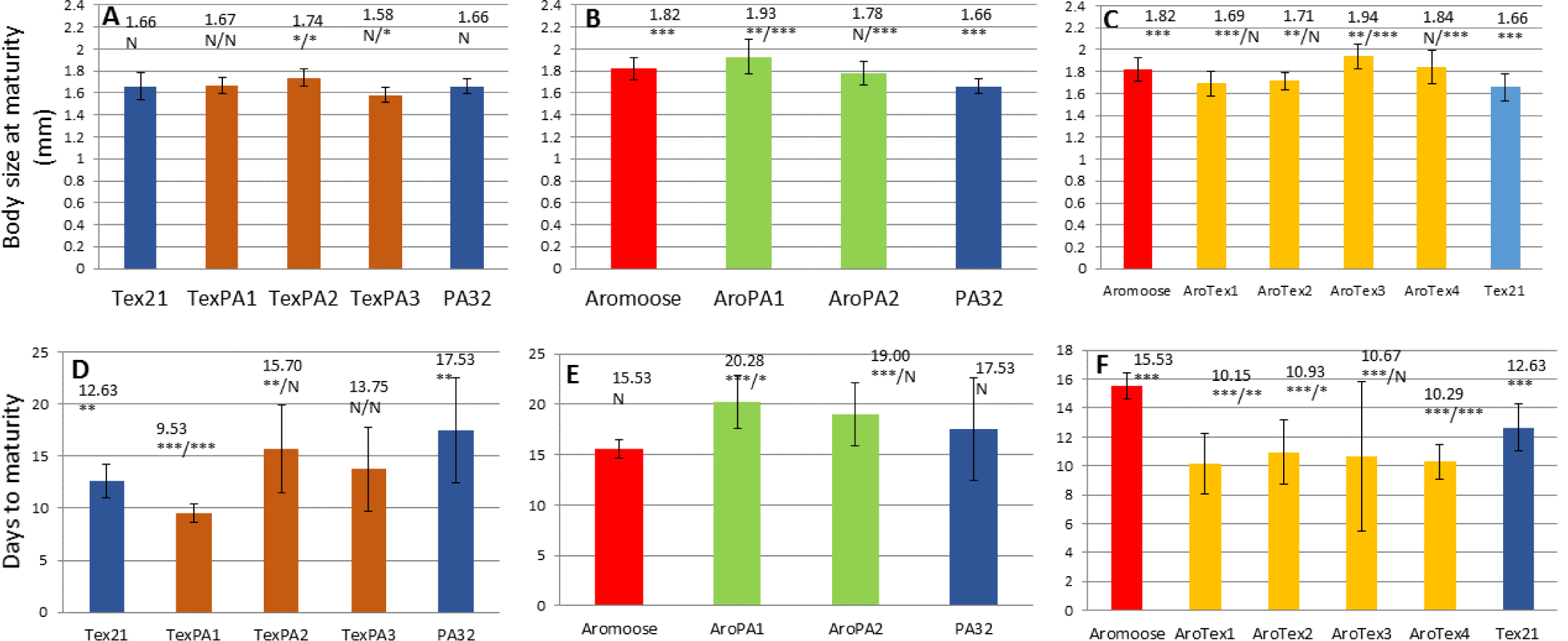
Female body size at maturity(A-C) and days to maturity (D-F) for parental isolates and their F_1_ crosses. Each panel represents results from one experimental cross. (A and D) males of Tex21 (blue) crossed with females of PA32 (blue) with 3 F_1_s (brown). (B and E) males of Aromoose crossed with females of PA32 with 2 F_1_s (green). (C and F) males of Aromoose crossed with females of Tex21 with 4 F_1_s (yellow).Above each bar, the numeric value of the trait is indicated. Below the numeric value, the significance of t-test is indicated. For the parental isolates the significance for comparing two parental clones is given, whereas forF_1_s the significance for comparing with paternal and maternal isolate is given respectively (divided by a forward slash). N: not significant; *: p < 0.05; **: p<0.01; ***: p<0.001.

Heterosis was observed in the F_1_s of all three crossing experiments, whereas no hybrid depression was found for this trait (**Figure 1A-C**). For our intra-specific *D. pulex* crosses (males of Tex21 crossed with females of PA32), the F_1_ hybrid TexPA2 was significantly bigger than both parents, averaging at 1.73 ±0.08 mm (*p*=0.005, *p*=0.04). TexPA1 showed no significant difference in size relative to both parents, averaging at 1.67 ±0.07 mm (*p*=0.24, *p*= 0.32). Lastly, TexPA3 (1.57 ±0.06 mm) was significantly smaller compared to PA32 (*p*= 0.02), but not significantly different from Tex21 (*p*=0.1).

When we compared the AroPA clones to their respective parents PA32 and AroMoose (**Figure 1B**), we found that body size at maturity for AroPA1 (1.92 ±0.15 mm) was significantly bigger than either parents (*p*=7.47×10^−7^, *p*=0.01). On the other hand, AroPA2 (1.78 ±0.10mm) was also significantly bigger than PA32 (*p*=0.001) but showed no significant difference relative to AroMoose (*p*=0.22).

When we examined the hybrids of AroMoose and Tex21, AroTex3 was significantly bigger compared to either AroMoose or Tex21, averaging at 1.93 ±0.11 mm (*p*=0.005, *p*=5.39×10^−6^). We also found that both AroTex1 and AroTex2 were significantly smaller than AroMoose but showed no significant difference relative to Tex21, with mean body size of 1.69±0.1 mm (*p*=0.001,*p*= 0.23) and 1.71±0.07 mm (*p*=0.002, *p*= 0.08), respectively. Lastly, AroTex4 (1.84 ±0.15 mm) was significantly bigger than Tex21 (*p*=0.001) but was not significantly different from AroMoose (*p*=0.31).

### Days to Maturity

Regarding the parental isolates (**Figure 1D-F**), the *D. pulicaria* isolate Aromoose (15.53 ±0.91 days) tends to mature faster than the *D. pulex* isolate PA32 (17.53± 5.05days), but no statistical significance was found. On the other hand, AroMoose matured significantly slower (p<0.01) than Tex21 that matured at 12.63 ±1.62 days, whereas Tex21 reached maturity significantly faster than PA32 (p<0.01).

One F_1_ from the intra-specific cross between Tex21 and PA32 showed sign of heterosis (**Figure 1D**), and 3 out of the 4 F_1_s in the inter-specific cross between Aromoose and Tex21 showed heterosis (**Figure 1F**). On the other hand, one hybrid (AroPA1) between Aromoose and PA32 showed declined fitness compared to both parental isolates (**Figure 1E**).

Specifically, we found that TexPA1 took significantly fewer days to mature compared to either parents Tex21 or PA32 at 9.53 ±0.91 days (*p*=1.07×10^−6^, *p*=8.41×10^−7^, **Figure 1D**). On the other hand, TexPA2 took significantly more days to mature compared to Tex21 (*p*=0.017) but showed no significant difference relative to PA32 (*p*=0.17), maturing at 15.70±4.16 days (**Figure 1D**). Lastly, TexPA3 matured at 13.75 ±4.03 days showed no significant difference when compared to both PA32 and Tex21 (*p*= 0.09, *p*=0.22, **Figure 1D**).

For hybrids between Aromoose and PA32 (**Figure 1E**), we found that AroPA1 took significantly more days to mature compared to both parents at 20.28 ±2.64 days (*p*=2.45×10^−7^, *p*=0.04). Interestingly, AroPA2 took significantly more days to mature when compared to Aromoose (*p*=3.53×10^−4^) but showed no significant difference when compared to PA32 (*p*=0.24), maturing at 19 ±3.16 days.

When we compared the AroTex clones to their parents Aromoose and Tex21 (**Figure 1F**), we found that AroTex1, AroTex2 and AroTex4 took significantly fewer days to mature when compared to both parents, maturing at 10.15 ±2.07 days (*p*=7.45×10^−10^, *p*= 0.002), 10.92 ±2.23 days (*p*=3.27×10^−8^, *p*=0.022) and 10.28±1.2 days (*p*=1.21×10^−13^, *p*=1.93×10^−4^), respectively. Moreover, AroTex3 took significantly fewer days to mature when compared to Aromoose (*p*=7.05×10^−4^) but was no different from Tex21 (*p*=0.12), maturing at 10.66 ±5.17days.

### Number of Broods

Among the parental isolates (**Figure 2A-C**), Tex21 showed the highest fecundity, with 11.45 ±3.95 broods during life span, which is significantly higher (*p*<0.05 in both tests) than both Aromoose (6.86 ±5.13) and PA32 (7.93 ±4.36 broods).We only identified one clear example of heterosis in the inter-specific hybrid AroPA2 (Aromoose crossed with PA32).

When we compared the TexPA clones to their respective parents Tex21 and PA32 (**Figure 2A**), we found that both TexPA1 (7.73 ±5.18 broods) and TexPA3 (4.25 ±2.5 broods) had significantly fewer broods when compared to Tex21 (*p*= 0.029, *p*=0.002), but showed no significant difference when compared to PA32 (*p*=0.45, *p*=0.06). Interestingly, TexPA2 showed no significant difference when compared to both parents, producing 10.1 ±5.74 broods (*p*=0.26, *p*=0.14).

**Figure 2.**
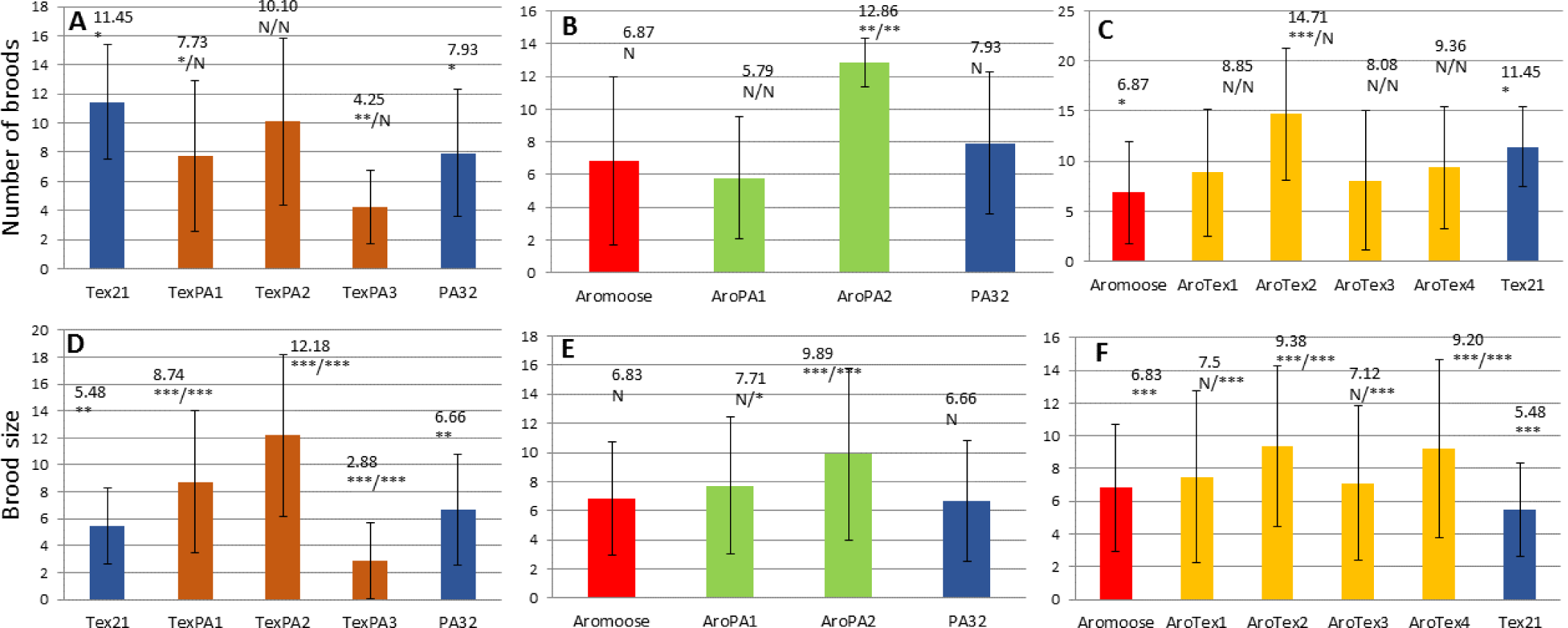
Number of broods (A-C) and brood size (D-F) for parental isolates and their F_1_ crosses. Each panel represents results from one experimental cross. (A and D) males of Tex21 (blue) crossed with females of PA32 (blue) with 3 F_1_s (brown). (B and E) males of Aromoose crossed with females of PA32 with 2 F_1_s (green). (C and F) males of Aromoose crossed with females of Tex21 with 4 F_1_s (yellow). Above each bar, the numeric value of the trait is indicated. Below the numeric value, the significance of t-test is indicated. For the parental isolates the significance for comparing two parental clones is given, whereas for F_1_s the significance for comparing with paternal and maternal isolate is given respectively (divided by a forward slash). N: not significant; *: p < 0.05; **: p<0.01; ***: p<0.001.

**Figure 3.**
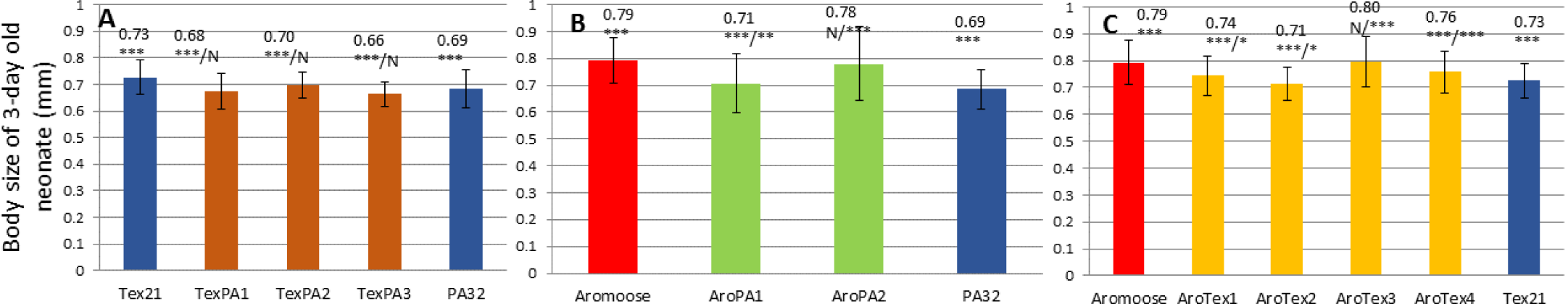
Body size of 3-day old neonates for parental isolates and their F_1_ crosses. Each panel represents results from one experimental cross. (A and D) males of Tex21 (blue) crossed with females of PA32 (blue) with 3 F_1_s (brown). (B and E) males of Aromoose crossed with females of PA32 with 2 F_1_s (green). (C and F) males of Aromoose crossed with females of Tex21 with 4 F_1_s (yellow). Above each bar, the numeric value of the trait is indicated. Below the numeric value, the significance of t-test is indicated. For the parental isolates the significance for comparing two parental clones is given, whereas for F_1_s the significance for comparing with paternal and maternal isolate is given respectively (divided by a forward slash). N: not significant; *: p < 0.05; **: p<0.01; ***: p<0.001.

When we compared the AroPA clones to their respective parents Aromoose and PA32 (**Figure 2B**), we found that AroPA1 showed no significant difference when compared to both parents, producing 5.78 ±3.72 broods (*p*= 0.26, *p*=0.08). On the other hand, AroPA2 produced significantly more broods when compared to both parents at 12.85 ±1.46 broods (*p*=0.003, p=0.004).

When we compared the AroTex clones to their parental isolates (**Figure 2C**), AroTex1, AroTex3 and AroTex4 showed no significant differences when compared to both parents, producing 8.84 ±6.36 broods (*p*=0.18, *p*= 0.12), 8.08 ±6.96broods (*p*= 0.3, *p* = 0.08) and 9.35±6.09 broods (*p* = 0.12, *p*=0.16), respectively. While showing no significant difference from Tex21 (*p*= 0.08), AroTex2 had significantly more broods comparing to Aromoose (*p*=6.36×10^−4^), producing 14.71±6.56 broods.

### Brood Size

Regarding the parental isolates (**Figure 2D-F**), the *D. pulex* isolate Tex21 (5.47±2.84 embryos per brood) appeared to have the lowest brood size, significantly lower than *D. pulex* isolate PA32 (6.65±4.12) and the *D. pulicaria* isolate Aromoose (6.82±2.87). No significant difference was detected between PA32 and Aromoose. For this trait, we identified heterosis in both intra- and inter-specific crossings, whereas only one F_1_ hybrid (TexPA3) had lower number of brood than both parents.

When we compared the TexPA clones to their respective parents Tex21 and PA32 (**Figure 2D**), we found that both TexPA1 and TexPA2 had significantly larger broods when compared to both parents, carrying 8.74 ±5.27 embryos (*p*=5.21×10^−11^, *p*=4.87×10^−4^) and 12.18 ±6.02 embryos (, *p*=1.55×10^−28^, *p*=3.61×10^−14^) per brood, respectively. On the other hand, TexPA3 had significantly smaller broods when compared to both parents, carrying 2.88 ±2.84 embryos (*p*=2.13×10^−4^, *p*=1.93×10^−4^) per brood.

When we examined the hybrids of AroMoose and PA32 (**Figure 2E**), we found that AroPA1 carried significantly larger broods when compared to PA32 (*p*=0.04), but showed no significant differences when compared to AroMoose (*p*=0.08), carrying on average 7.70 ±4.71 embryos per brood. However, AroPA2 had carried significantly larger broods when compared to both parents, carrying 9.88±5.87 embryos (*p*=1.35×10^−5^, *p*=3.5×10^−6^) per brood.

When we compared the AroTex clones to their respective parents AroMoose and Tex21 (**Figure 2F**), we found that both AroTex1 and AroTex3 carried significantly larger broods when compared to Tex21 (*p*=1.97×10^−5^, *p*=2.54×10^−4^), but showed no significant difference when compared to AroMoose (*p*= 0.14, *p*= 0.31), carrying on average 7.5 ±5.25 embryos and 7.11 ±4.71 embryos per brood, respectively. However, both AroTex2 and AroTex4 carried significantly larger broods when compared to both parents, carrying 9.37 ±4.89 embryos (*p*=2.84×10^−6^, *p*=1.67×10^−18^) and 9.20 ±5.44 embryos (*p*=1.18×10^−4^, *p*=1.61×10^−13^) per brood, respectively.

### Body size of 3-day old neonate

Among the parental isolates (**Figure 5**), the *D. pulicaria* isolate AroMoose has the largest body size for 3-day old neonates (0.79±0.08 mm), significantly larger (p<0.001 in both cases) than both *D. pulex* isolates PA32 (0.69±0.07mm) and Tex21 (0.73±0.06mm). We noted that the difference between PA32 and Tex21 was also statistically significant (p<0.001). It is worth noting that no examples of heterosis was identified in any of the F1 hybrids of intra- and inter-specific crosses.

When we compared the TexPA clones to their respective parents PA32 and Tex21, we found that all TexPA clones, TexPA1 (0.67±0.06mm) and TexPA2 (0.70±0.04mm) and Tex3, averaging at 0.66±0.04mm were significantly smaller when compared to Tex21 (*p*=8.15×10^−14^, *p*=9.37×10^−7^, *p*=6.78×10^−6^, respectively), but showed no significant differences when compared to PA32 (*p*= 0.07, *p*= 0.05, *p*= 0.08, respectively).

When examining hybrids of Aromoose and PA32, we found that AroPA1 was significantly larger when compared to PA32 (*p*= 0.01), and significantly smaller when compared to Aromoose (*p*=2.7×10^−13^), averaging at 0.71±0.1mm. AroPA2 was also significantly larger when compared to PA32 (*p*=2.04×10^−13^) but showed no significant difference when compared to Aromoose (*p*=0.17), averaging at 0.78±0.13 mm.

When comparing the AroTex clones with their respective parents AroMoose and Tex21, we found that both AroTex1 and AroTex4 were significantly larger when compared to Tex21 (*p*= 0.02, *p*=2.05×10^−5^), but significantly smaller when compared to AroMoose (*p*=2.45×10^−6^, *p*=7.21×10^−5^), averaging at 0.74± 0.07 mm and 0.76± 0.07mm, respectively. We also found that AroTex2 was significantly smaller when compared to both parents, averaging at 0.71±0.06 mm (*p*=2.63×10^−20^, *p*= 0.03). Lastly, AroTex3 was significantly larger when compared to Tex21 (*p*=3.26×10^−14^) but showed no significant difference when compared to AroMoose (*p*=0.32), averaging at 0.80± 0.09 mm.

## Discussion

In this study we examined the life-history traits of intra-specific hybrids of *D. pulex* and interspecific hybrids between *D. pulex* and *D. pulicaria*. Our motivation is to evaluate whether there are major genetic incompatibilities between these two species affecting life-history traits. As life-history traits can be easily affected by environmental conditions, we kept the experimental animals for two generations in constant environmental conditions prior to using the third-generation individuals for life history assay. This procedure eliminates the (grand)maternal effects to the maximal extent and the observed differences in life history traits are most likely a reflection of the true genetic differences among the examined *Daphnia* isolates.

Although the sample sizes of F_1_ hybrids for our intra- and inter-specific are low, we made a few important observations regarding the fitness outcome of hybridization. First, hybrids displaying heterosis re identified for all life-history traits examined except for body size of 3-day old neonates. Both intra- and inter-specific crosses produced hybrids with heterosis for female size at maturity, days to maturity, and brood size, whereas heterosis was only identified in one inter-specific cross (AroMoose x PA32) for the number of broods. This observation suggests that at both the intra- and inter-specific level there are no fixed genetic elements that result in decreased fitness in hybrids, although some hybrids also show declined fitness (see below). This observation is also consistent with the idea that *D. pulex* and *D. pulicaria* are at an early stage of speciation, where heterosis can still occur. As to why the heterosis occurs for these life-history traits, current theories suggest several possible explanations, e.g., uniparentally inherited elements (Fraisse et al., 2016), or breakdown of deleterious epistatic interaction in the parental population (Dagilis et al., 2019). As *Daphnia* populations can experience inbreeding (Lynch et al., 2017), within each pond or lake population deleterious mutations could accumulate due to the reduced effective population size and reduced power of natural selection. We argue that the breakdown of negative epistasis interaction is likely a cause of heterosis that we observed in these experiments.

A segregation and recombination in sexual reproduction breaks down the parental intra- and inter-locus association of alleles and create new combinations of alleles (Otto, 2009), the offspring could suffer declined fitness compared to either parental genotype. This is the so-called genetic slippage (Lynch & Deng, 1994). In our crossing experiments, we identified a few instances of F_1_s with inferior fitness relative to both parents, e.g., days to maturity (AroPA1) and brood size (TexPA3). Moreover, we observed some hybrids demonstrating significantly lower fitness relative to one parent but not different form the other parent, e.g., number of broods (TexPA1 and TexPA3 in **Figure 2**). Or some hybrids demonstrating intermediate values for traits relative to parents, which occurred for all traits across the board.

Our observations have several implications for understanding the divergence of *D. pulex* and *D. pulicaria*. As natural F_1_ hybrid populations between *D. pulex* and *D. pulicaria* are rarely found (Heier & Dudycha, 2009), it is hypothesized that F_1_ hybrids are selected against in natural habitat of pond and lakes. Our data show that the F_1_ hybrids can have great variation on life-history traits relative to the parental genotypes, e.g., heterosis, intermediate values, and declined fitness. If F_1_ hybrids occur in nature, it is likely some of the hybrids would survive regardless of the direction of selection and facilitate gene flow between these two species. As a matter of fact, introgression from *D. pulicaria* to *D. pulex* has been linked to the cause of two phenotypic changes, i.e., the origin of obligate asexual reproduction (Xu et al., 2015) and the suppression of male production (Ye et al., 2019). We note that most previous studies used a few characteristic molecular marker (e.g., allozyme locus LDH) to identify the populations (e.g., Hebert et al., 1989; Hebert & Crease, 1983). However, introgression in genomic regions other than these assayed loci may not be detected. With the increasingly easy access to high throughput sequencing, we suggest that it is necessary to scrutinize a population on the genome-wide scale to discern its identity.

Moreover, as the fitness of F_1_ hybrids does not seem to form a strong barrier for hybridization/introgression to occur, it is possible that F_1_ hybrids would quickly backcross with local resident population (e.g., with *D. pulex* population in the event of *D. pulicaria* migrates a to pond population). Therefore, F_1_ hybrids would be a transient phenomenon and would be hard to capture in nature. This scenario again reminds us about the importance of analyzing the ancestry of populations of interests from a genome-wide perspective as backcrossing would create large sections of the genome homozygous of alleles from the backcrossing parental species, which reduces the power of using a few diagnostic markers for ancestry identification. Similarly, selfing among F_1_s can create long tracks of homozygosity, which can also mislead ancestry identification based on a few markers. We therefore suggest that introgression between these two species may be common and may be detected by using genome-wide DNA sequence data.

## Acknowledgements

We thank Trung Huynh, Swatantra Neupane, Marelize Snyman, Duc Ly for their help with the experiments. This work is supported by NIH grant R35GM133730 to SX.

## Notes

### Competing Interest Statement

The authors have declared no competing interest.

